# Hearing the World Differently: Examining Predictive Coding Accounts of Autism Using MEG

**DOI:** 10.1101/2022.10.03.510718

**Authors:** Hannah Rapaport, Elizabeth Pellieano, Robert A. Seymour, Nicholas Benikos, Wei He, Yanan Sun, Jon Brock, Paul F. Sowman

## Abstract

Predictive coding accounts of autism suggest that autistic perception is characterised by divergent precision weighting. The precise nature of this divergence, however, is debated. Here, we sought to disentangle competing predictive coding accounts of autism by testing them at a neural level. To this end, we used paediatric magnetoencephalography to record the auditory evoked fields of 10 young autistic children (*M* = 6.2 years, range = 4.2– 8.6) and 63 neurotypical children (*M* = 6.1 years, range = 3.0– 9.8) as they listened to a roving auditory oddball paradigm. For each participant, we subtracted the evoked responses to the ‘standard’ from the ‘deviant’ pure tones to calculate the mismatch field ‘MMF’: an electrophysiological component that is widely interpreted as a neural signature of predictive coding. We found no significant differences between the two groups’ MMF amplitudes, *p* > .05. An exploratory analysis indicated larger MMF amplitudes in most of the autistic children compared to their average-age-matched neurotypical counterparts, *p* < .05. We interpret these findings as preliminary evidence in support of the ‘inflexibly high prior and sensory precision’ account, and against the ‘inflexibly low prior-relative-to-sensory precision’ accounts of autistic perception.

**Highlights:**

- We used paediatric MEG to compare autistic and neurotypical MMFs amplitudes.
- Exploratory case-cohort analyses revealed mostly larger MMFs in autistic cases.
- Larger MMFs support the notion of precise, inflexible prediction errors in autism.

## 1. Introduction

Autism is a lifelong, heritable and heterogeneous neurodevelopmental condition that shapes how a person experiences and interacts with their environment (Lord et al., 2020). According to the DSM-5, autism is characterised by differences in social interaction and communication (e.g., difficulties with social reciprocity, nonverbal communication, and social relationships), as well as so-called ‘restricted, repetitive patterns of behaviour, or interests or activities’ (e.g., repetitive motor movements, or ‘stimming’ behaviours, a strong desire for sameness, intense interests, and, most recently, differences in sensory perception; American Psychiatric Association, 2013).

Whereas earlier ‘coherence’ accounts of autism (Frith, 1989; Frith & Happé, 1994; Happé & Booth, 2008; Happé & Frith, 2006; Mottron et al., 2006; Mottron & Burrack, 2001) struggled to provide a comprehensive account of autistic differences—particularly with regards to hypo-sensitivities and ‘restricted and repetitive behaviours’, more recent ‘predictive coding’ accounts (Brock, 2012; Friston et al., 2013; Lawson et al., 2014, 2017; Pellicano, 2013; Pellicano & Burr, 2012; van Boxtel & Lu, 2013; Van de Cruys et al., 2013, 2014) hold promise in offering a unifying account of autistic differences across the social and non-social domains. Here we sought to disentangle competing predictive coding accounts of autism by testing them at a neural level, using paediatric magnetoencephalography (MEG) in conjunction with a roving auditory oddball paradigm.

### 1.1. Predictive Coding Accounts of Autism

Under predictive coding, the brain embodies an internal, probabilistic, generative model which represents the statistical structure of the physical world (Clark, 2015; Hohwy, 2013; Rao & Ballard, 1999) and uses it to generate predictions about the most likely (hidden) physical causes of incoming sensory signals. The brain then tests its model in a hierarchical fashion, comparing its top-down prediction (generated in higher brain regions) against the bottom-up sensory signal (propagated up the system from lower brain regions that are closer to the sensory peripheries). The prediction and sensory signal can be represented as probability distributions and the difference between the distributions the prediction error (PE). PE represents the information the model failed to predict and which requires further processing at higher levels of the neural hierarchy. The central tenet of predictive coding theory is that the brain strives to reduce processing requirements by minimising PEs over time (Clark, 2013).

Divergent precision weighting has been proposed to account for unique autistic perception. Specifically, some predictive coding accounts of autism suggest that autistic perceptual and behavioural differences may be explained by a system that maintains either ‘inflexibly low prior precision’ (Pellicano, 2013; Pellicano & Burr, 2012) or, conversely, ‘inflexibly high sensory precision’ (Brock, 2012)—both of which would result in a perceptual experience that is less biased by prior knowledge and more biased towards incoming sensory signals.

Alternatively, a third account has proposed ‘inflexibly high prior and sensory precision’ in autistic people (Van de Cruys et al., 2013, 2014). In a stable environment, attempting to match a precise ‘prior’ distribution to a precise and invariable ‘sensory signal’ would give rise to very low-precision PEs. Yet, in a volatile environment, attempting to match a precise ‘prior’ distribution to a precise and variable ‘sensory signal’ would give rise to high-precision PEs, motivating constant model updating and learning (hence, the name of the account: “High, Inflexible Precision of Prediction Errors in Autism”; HIPPEA).

A fourth account has proposed ‘inflexibly low prior and sensory precision’ (Lawson et al., 2017). It is difficult, however, to reconcile this account with the core characteristics of autism. For example, the suggestion that autistic adults are “less surprised than neurotypical adults when expectations are violated” (Lawson et al., 2017, p. 1293) is at odds with the evidence that autistic people often have difficulties coping with unpredictability of everyday life (Simmons et al., 2009). Furthermore, this account does not explain the occurrence of stimming or self-injurious behaviours in autistic people.

### 1.2. Testing the Predictive Coding Accounts of Autism

A popular approach for testing predictive coding is to use electrophysiological techniques (e.g., MEG or electroencephalography; EEG) to record participant’s brain responses as they listen to an auditory oddball paradigm (Friston, 2005; Heilbron & Chait, 2017). These paradigms are comprised of auditory stimuli (e.g., pure tones or speech sounds) classed as either high-probability ‘standards’ or lower-probability ‘deviants’. In adults, averaged evoked responses to the ‘standards’ and ‘deviants’ typically diverge between 100 and 250 ms following stimulus onset, with the ‘deviant’ waveform showing a larger amplitude relative to the ‘standard’ waveform (Näätänen et al., 2019). This divergence, which is conventionally presented in a ‘deviant-minus-standard’ difference waveform, is the ‘mismatch negativity’ (MMN, EEG literature, or the ‘mismatch field’ (MMF, MEG literature; hereafter, the MMN/F). The MMN/F is widely considered to be a neural index of PE (Friston, 2005), thought to reflect a larger PE to the lower-probability ‘deviants’ relative to the higher-probability ‘standards’. This interpretation is based on evidence that the evoked response amplitude increases as an inverse function of stimulus probability (Denham & Winkler, 2020).

We tested several competing hypotheses regarding how predictive brain function (as indexed by the MMF) might differ between autistic and neurotypical children. Under the inflexibly low prior-relative-to-sensory precision accounts (Brock, 2012; Pellicano & Burr, 2012), autistic children’s brains may be relatively less proficient at extracting statistical regularities from the auditory sequence compared to their neurotypical counterparts. Thus, autistic children may form relatively less-precise predictions for the upcoming stimuli. This, in turn, would give rise to similar magnitudes of PE for the high-probability standards and lower-probability deviants, as indexed by a relatively attenuated MMF amplitude in autistic children.

Conversely, under the HIPPEA account (Van de Cruys et al., 2014), the pseudorandom deviant stimuli may elicit unduly precise PE signals, motivating the brain to update its model of the environment based on the unpredicted information. Thus, all subsequent repetitions of the ‘deviants’ (which, through repetition, become ‘standards’) would be precisely predicted, thereby eliciting minimal PE signals. Overall, autistic children’s brains may generate larger PE signals (i.e., evoked responses) to ‘deviants’ relative to ‘standards’, resulting in a relatively large MMF amplitude.

Yet, existing MMN/F findings do not provide clear guidance as to which of these two hypotheses might be more likely. Perhaps the most robust evidence to date comes from a systematic review and meta-analysis (Schwartz et al., 2018), which reported that young autistic children had attenuated MMN/F amplitudes compared to their neurotypical counterparts. Furthermore, findings from a large number of individual studies are mixed, with evidence of autistic children showing MMN/F amplitudes that are smaller (Andersson et al., 2013; Dunn et al., 2008; Jansson-Verkasalo et al., 2005; Ludlow et al., 2014; Seri et al., 1999; Yoshimura et al., 2017), larger (Ferri et al., 2003), or not significantly different from that of their neurotypical counterparts (Gomot et al., 2002, 2011; Lepistö et al., 2009; Roberts et al., 2011; Weismuller et al., 2015). Still others have reported mixed within-study findings related to the type of stimulus employed and the selection of sensors analysed (Jansson-Verkasalo et al., 2003; Korpilahti et al., 2007; Kujala et al., 2010; Lepistö et al., 2005, 2006, 2008; Yu et al., 2015).

Finally, most previous MMN/F studies recruited autistic and neurotypical children aged six years and over, whereas those in the pre-school years (i.e., between the ages of three and five years; Centers for Disease Control and Prevention, 2021) have received far less empirical attention. Investigating the MMF amplitude in autistic and neurotypical pre-school children is of particular interest, as findings could indicate a divergence in predictive brain function at the stage of life in which most children receive an autism diagnosis (Gibbs et al., 2019).

### 1.3. The Current Study

We sought to disentangle the competing predictive coding accounts of autism. For this purpose, we used paediatric MEG in conjunction with a ‘roving’ auditory oddball paradigm and compared MMF amplitudes between autistic and neurotypical children, aged between three and eight years. Attenuated MMF amplitudes in autistic children, compared to their neurotypical counterparts, would be consistent with the ‘inflexibly low prior-relative-to-sensory precision’ accounts (Brock, 2012; Pellicano, 2013; Pellicano & Burr, 2012). Conversely, relatively larger MMF amplitudes in autistic children would be consistent with the ‘HIPPEA’ account (Van de Cruys et al., 2013, 2014).

## 2. Methods

### 2.1. Ethical Approval

All procedures were approved by the Macquarie University Human Research Ethics Committee (reference numbers: 5201300834 and 5201600188). Informed consent was obtained from participants’ caregivers and, where possible, from the children themselves. All participants received 40 AUD and a gift bag for their participation. Autistic participants received an additional 20 AUD to compensate for the extra time spent undergoing the ADOS-2 assessment.

### 2.2. Participants

#### 2.2.1. Autistic Children

Twenty-one children (aged 3 to 8 years) who had an independent, clinical diagnosis of autism were recruited via advertisements posted on social media, and flyers distributed to local schools, autism-focused child-care centres, medical clinics, and autism service providers. Of the 21 recruited children, 11 were unable to complete the testing session due to: (i) a sensory aversion to some aspect of the testing environment (*n* = 10) – including the MEG cap (*n* = 6; *M*_age_ = 4.2 years, range: 3.0 – 7.8), the MEG dewar (*n* = 2; *M*_age_ = 5.8 years), the auditory oddball stimuli (*n* = 1; aged 3.4 years), and the digitiser pen (*n* = 1; aged 5.0 years) – or (ii) having outgrown the child MEG system dewar (*n* = 1; aged 8.3 years; see Supplementary Table S1 for further details).

The final sample therefore consisted of 10 autistic children (*M*_age_ = 6.2 years, range = 4.2–8.6 years; see Table 1). All had normal, or corrected-to-normal, vision and had no history of epilepsy or brain injury, as reported by parents. We found no significant differences between the ‘included’ and ‘excluded’ autistic children with respect to age, ADOS-2 Comparison Scores, or SCQ Lifetime scores (see Table 2). However, the mean age was lower for the excluded children, and the effect size was high. Furthermore, the two groups appeared to differ in their spoken language ability: nine of the 10 included children (90%) obtained the lowest rating for ADOS-2 assessment Item A1 (‘Overall Level of Non-Echoed Spoken Language’), indicating *more* complex spoken language, compared to only four of the 11 excluded children (36%). As such, it seems that the ‘included’ children were older and had more complex expressive language.

**Table 1.** Demographic Details of the Autistic Children and their Average-Age-Matched Neurotypical Counterparts, and Latencies of Significant Case-Cohort MMF Amplitude Differences

| Autistic children |  |  |  |  |  |  | Neurotypical average-age-matched comparison group |  | Autistic-neurotypical comparison |  |
| --- | --- | --- | --- | --- | --- | --- | --- | --- | --- | --- |
| ID | ADOS-2 |  |  | SCQ | Sex | Age | Average age | N | Latencies (s) of sig. MMF amplitude diff. |  |
|  | CSS | Module | Item A1 |  |  |  |  |  | Start | End |
| P1 | 9 | 1 | Complex | 19 | F | 4.2 | 4.2 | 17 | N/A | N/A |
| P2 | 3 | 2 | Complex | 17 | F | 4.5 | 4.5 | 20 | 0.03 | 0.1 |
| P3 | 9 | 1 | More than five<br>single words | 14 | M | 4.6 | 4.6 | 20 | 0.15 | 0.22 |
|  |  |  |  |  |  |  |  |  | 0.23 | 0.33 |
| P4 | 4 | 2 | Complex | 25 | F | 4.8 | 4.8 | 24 | N/A | N/A |
| P5 | 8 | 2 | Complex | 27 | M | 5.8 | 5.8 | 32 | 0.05 | 0.12 |
| P6 | 10 | 3 | Complex | 17 | M | 6.8 | 6.8 | 31 | 0.2 | 0.4 |
| P7 | 10 | 3 | Complex | 19 | M | 7.2 | 7.2 | 32 | 0.05 | 0.14 |
| P8 | 10 | 3 | Complex | 9 | M | 7.2 | 7.2 | 32 | 0.12 | 0.19 |
|  |  |  |  |  |  |  |  |  | 0.32 | 0.39 |
| P9 | 10 | 3 | Complex | 11 | M | 8.0 | 8.0 | 13 | 0.2 | 0.31 |
| P10 | 10 | 3 | Complex | 24 | M | 8.6 | 8.6 | 10 | 0.0 | 0.08 |
|  |  |  |  |  |  |  |  |  | 0.27 | 0.36 |

Notes. P = Participant. ADOS-2 = Autism Diagnostic Observation Schedule – 2<sup>nd</sup> edition (Lord et al., 2012); CSS = Calibrated Severity Score (Gotham et al., 2009), maximum = 10. Item A1 = Overall Level of Non-Echoed Spoken Language. SCQ = Social Communication Questionnaire (Rutter et al., 2003) score. Higher scores on the ADOS-2 and SCQ reflect greater autistic severity.

**Table 2.** Descriptive Statistics and Group Differences for Age, ADOS-2, and SCQ Scores for the Included and Excluded Autistic Participants

|  | *n* | *M* | *SD* | Min | Max | *t* | *p* | Cohen's *d* |
| --- | --- | --- | --- | --- | --- | --- | --- | --- |
| Age (years) |  |  |  |  |  | 1.72 | .102 | 0.75 |
| Included | 10 | 6.16 | 1.60 | 4.17 | 8.58 |  |  |  |
| Excluded | 11 | 4.86 | 1.86 | 3.00 | 8.25 |  |  |  |
| ADOS-2 |  |  |  |  |  | 1.41 | .174 | 0.62 |
| Included | 10 | 8.30 | 2.63 | 3 | 10 |  |  |  |
| Excluded | 11 | 6.82 | 2.18 | 3 | 9 |  |  |  |
| SCQ |  |  |  |  |  | 0.68 | .946 | 0.03 |
| Included | 10 | 18.20 | 5.92 | 9 | 27 |  |  |  |
| Excluded | 9<sup>a</sup> | 18.00 | 6.86 | 4 | 27 |  |  |  |

*Notes.* ADOS-2 = Autism Diagnostic Observation Schedule – 2nd edition (Lord et al., 2012) calibrated severity score (Gotham et al., 2009), maximum = 10. SCQ = Social Communication Questionnaire (Rutter et al., 2003) score. Higher scores on the ADOS-2 and SCQ reflect greater autistic severity. <sup>a</sup>Two datasets were missing as some parents did not complete this form.

#### 2.2.2. Neurotypical Children

The neurotypical children were recruited as part of a separate, two-phase study via both the Macquarie University ‘Neuronauts’ child research participation database and an advertisement in the ‘Sydney’s Child’ magazine. The two testing phases (which were separated by 20 months, on average) produced 32 and 31 high-quality datasets, respectively. Combining these samples resulted in 63 neurotypical datasets (*M*_age_ = 6.1 years, range = 3.0–9.8 years). These 63 neurotypical children served as comparisons to the autistic children. All neurotypical children had normal or corrected-to-normal vision and had no history of developmental disorders, epilepsy, brain injury or language or speech impairment, as reported by parents.

### 2.3. Behavioural Assessments

The Autism Diagnostic Observation Schedule – 2^nd^ edition (ADOS-2; Lord et al., 2012) was used to assess current autistic features for the autistic participants only. The ADOS-2 is a 40-minute, standardised observational scale, designed to present opportunities for the evaluation of social, communicative, and repetitive behaviours. Raw algorithm scores were converted to standardised ADOS-2 calibrated severity scores (Gotham et al., 2009), as these scores are less affected by factors such as age, language, and cognitive ability. Severity scores range from 1 to 10 where higher scores reflect greater autistic severity.

The Lifetime version of the Social Communication Questionnaire (SCQ; Rutter et al., 2003) was completed by the parents of the autistic participants and used to index the degree of autistic features. The SCQ’s 40 items are derived from the well-validated Autism Diagnostic Interview (ADI; Le Couteur et al., 1989) with which it has good agreement (Corsello et al., 2007). Scores range from 0 to 39, where higher scores indicate greater social communication difficulties. Scores of 15 or more are considered indicative of clinically significant autistic features.

### 2.4. Auditory Oddball Paradigm

Electrophysiological responses were measured as participants listened to a passive ‘roving’ auditory oddball paradigm (Garrido et al., 2008; see Figure 1). Pure sinusoidal tones (duration: 70 ms, including a 5 ms rise and fall times; inter-stimulus interval: 500 ms; average volume: 80 SPL) were presented in sequences consisting of 1 to 7 stimuli. Each sequence differed from the previous sequence in pitch (500–800 Hz in steps of 50 Hz). The length and pitch of the sequence varied pseudo-randomly. The first and the last tone in each sequence were defined as the ‘deviant’ and ‘standard’ stimuli, respectively. Evidently, the ‘standards’ and ‘deviants’ in each sequence had identical physical properties, differing only in their order of presentation. The paradigm ran for 15 minutes. No instructions were required. The paradigm was programmed and presented in MATLAB (Mathworks, Natick, MA, USA) using Cogent 2000 version 1.32 (http://www.vislab.ucl.ac.uk/cogent.php).

**Figure 1.**
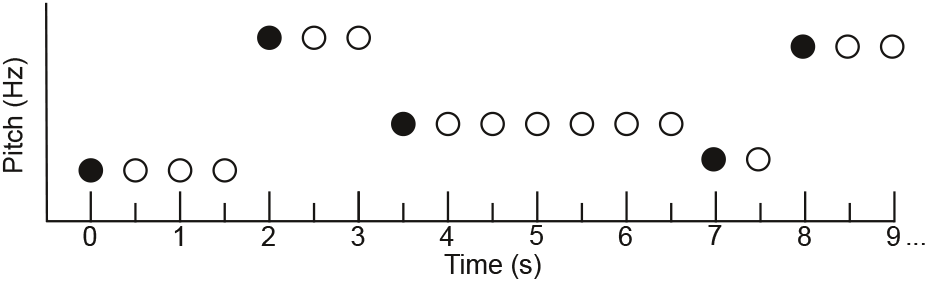
A schematic illustration of the roving auditory oddball paradigm (Garrido et al., 2008). The first tone in each sequence is a ‘deviant’ (black circles) and the final tone in each sequence is the ‘standard’ (or the ‘pre-deviant; the white circle prior to the black circle).

### 2.5. MEG Acquisition

The electrophysiological data were acquired using a whole-head, supine, paediatric MEG system (Model PQ1064R-N2m, KIT, Kanazawa, Japan), housed in a magnetically shielded room (MSR; Fujihara Co. Ltd., Tokyo, Japan). The MEG sensor array consisted of 125 first-order axial gradiometers, each of which had a coil diameter of 15.5 mm and a baseline of 50 mm (for further details, see He et al., 2019). The dewar was designed to fit a maximum head circumference of 53.4 cm, accommodating the heads of more than 90% of 5-year-old Caucasian children (Johnson et al., 2010). All data were acquired at a sampling rate of 1000 Hz and with an on-line bandpass filter of 0.03–200 Hz. To maximise the likelihood of obtaining high-quality data, we followed a child-friendly MEG testing protocol (which is available as an open-access video article; see Rapaport et al., 2019).

Before the MEG recording, participants were fitted with a polyester cap containing five head position indicator (HPI) ‘marker’ coils. A digitiser pen (Polhemus Fastrak, Colchester, USA) was used to record the locations of the HPI coils, as well as three fiducial points (the nasion and bilateral pre-auricular points) and 300-500 points from the scalp and face. For children who could not tolerate the Polhemus procedure, we instead used a contact-free 3D iPad head scan (Seymour, 2019; https://bit.ly/3alPGvl). The iPad procedure involved the use of a ‘Scanner–Structure SDK’ iPad application (Occipital, Inc., Boulder, CO)and a Structure Sensor accessory (Occipital Inc., Boulder, CO), which was mounted onto the iPad tablet camera lens.

During the MEG recording, children listened to the auditory oddball paradigm whilst watching a silent video of their choice. The paradigm was presented via a 60*60cm speaker (Panphonics SSH sound shower, Panphonics), which was positioned centrally at the foot of the plinth. The video was projected onto the ceiling of the MSR above the dewar. Participants’ head position in relation to the MEG sensor array was continuously monitored using a real-time marker coil tracking system (Oyama et al., 2012). Children were accompanied by a researcher (and, most often, their caregiver) for the duration of the MEG recording. Following the MEG recording, autistic children completed the ADOS-2 and their parents the SCQ. The entire session ran for approximately 45 minutes for the neurotypical children and 90 minutes for the autistic children.

### 2.6. Data Pre-processing

The following steps were performed in MEG160 (Yokogawa Electric Corporation and Eagle Technology Corporation, Tokyo, Japan). Channels that were consistently saturated for more than 10% of the recording were excluded from the subsequent steps on an individual basis. Environmental noise—estimated based on recordings from three reference magnetometers—was suppressed using a Time-Shift Principal Component Analysis (TSPCA) algorithm (de Cheveigné & Simon, 2007; block width: 10,000 ms, 3 shifts). Data acquired with the real-time marker coil tracking system were used to correct for head motion artefacts (Knösche, 2002; realignment conditions: sphere mesh = 321, prune ratio = 0.05).

Further pre-processing steps were performed in Matlab 2020a (MathWorths, Inc., Natick, MA, USA) using the Fieldtrip Toolbox v20200213 (Oostenveld et al., 2011). For each participant, the entire recording was high- and low-pass filtered at 0.1 and 40 Hz, respectively (using a onepass-zerophase firws filter with a Blackman window), and band-stop filtered to remove residual 50 Hz power-line contamination and its harmonics. Following visual inspection, segments of the recordings containing artefacts (e.g., SQUID jumps and jaw clenches) were removed. Channels that contained a large number of these visually-identified artefacts and/or were saturated for more than 10% of the recording were interpolated to ensure that all participants had the same number of MEG channels (Medvedovsky et al., 2007). An independent component analysis (ICA) was used to suppress eye blink artefacts. First, the raw recordings were high-pass filtered at 1Hz to improve the ICA performance (Winkler et al., 2015). Subsequently, up to two components with scalp distributions corresponding to eye blinks were removed from the 0.1 Hz, pre-processed data.

The continuous data were epoched into segments of 500 ms (100 ms pre- and 400 ms post-stimulus onset). Standard epochs were defined as the final trial in a sequence that consisted of more than two trials. Deviant epochs were defined as the first trial in a sequence that immediately followed a sequence consisting of more than two trials. There was an average of 268 deviant and 268 standard epochs for each autistic participant, and 266 deviant and 268 standard epochs for each neurotypical participant. The first deviant epoch was excluded from further analysis. All ‘standard’ and ‘deviant’ epochs were averaged across, respectively, to compute ‘standard’ and ‘deviant’ event-related fields (ERFs). ‘Standard’ and ‘deviant’ global field powers (GFPs) were calculated to quantify the amount of absolute activity at each time point across all of the sensors. Subtracting the ‘standard GFP’ from the ‘deviant GFP’ produced an ‘MMF^1^-GFP’ for each participant.

### 2.7. Statistical Analyses

The following statistical analyses were performed in MNE-Python (Gramfort et al., 2013). We compared MMF amplitudes between the autistic and neurotypical groups using a non-parametric cluster-based permutation analysis with independent-samples *t*-tests (Maris & Oostenveld, 2007). This approach has been shown to adequately control the Type-I error rate for electrophysiological data. We clustered samples whose *t*-values fell below a threshold corresponding to an alpha level of 0.05 (on the basis of temporal proximity) and calculated cluster-level test statistics by taking the average of the *t*-values within each cluster. The data were then permuted 1,000 times, each time randomly shuffling the group labels and recomputing the *t*-values. We constructed a permutation distribution from these random partition *t*-values. Finally, the significance of each cluster determined by using a threshold Monte-Carlo *p*-value.

We also conducted a series of exploratory case-cohort analyses using permutation tests. The following steps were performed separately for each of the 10 autistic children. First, we created an average-age-matched group of neurotypical children (whose ages were within 12 months of the autistic child), and subtracted the autistic child’s MMF from the MMF of each child in the neurotypical comparison group. We then used permutation one-sample *t*-tests to determine whether the difference in MMF amplitudes between the autistic child and the neurotypical comparison group was significantly different from zero (Maris & Oostenveld, 2007). We selected all samples whose *t*-values fell below a threshold corresponding to an alpha level of 0.01 and clustered the selected samples based on their temporal proximity. We then calculated cluster-level test statistics by taking the sum of the *t*-values within each cluster. Finally, the data were permuted 1,000 times, we constructed a permutation distribution and tested the significance of each cluster using a threshold Monte-Carlo *p*-value. To visualise where each autistic child fell in relation to their neurotypical counterparts, we computed mean MMF amplitudes (by averaging the individual data over each of the significant time windows) and plotted the individual data for each case-cohort comparison.

## 3. Results

### 3.1.1 Group-Level Comparison

**Figure 2** shows the ‘standard’ and ‘deviant’ GFPs and **Figure 3** shows the ‘MMF-GFPs’ for the autistic and neurotypical children, respectively. We found no significant differences between the two groups’ MMF amplitudes, *p* > .05.

**Figure 2.**
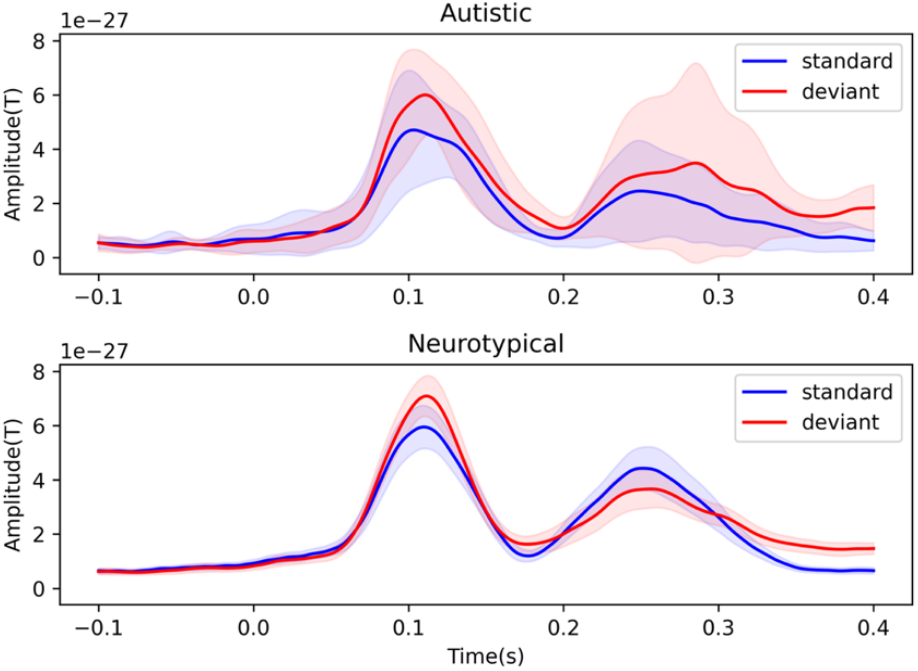
Group-level global field power (GFP) amplitudes (Tesla) by time (ms) for the ‘standard’ (blue) and ‘deviant’ (red) conditions. Top panel: autistic children (*N* = 10). Bottom panel: neurotypical children (*N* = 63).

**Figure 3.**
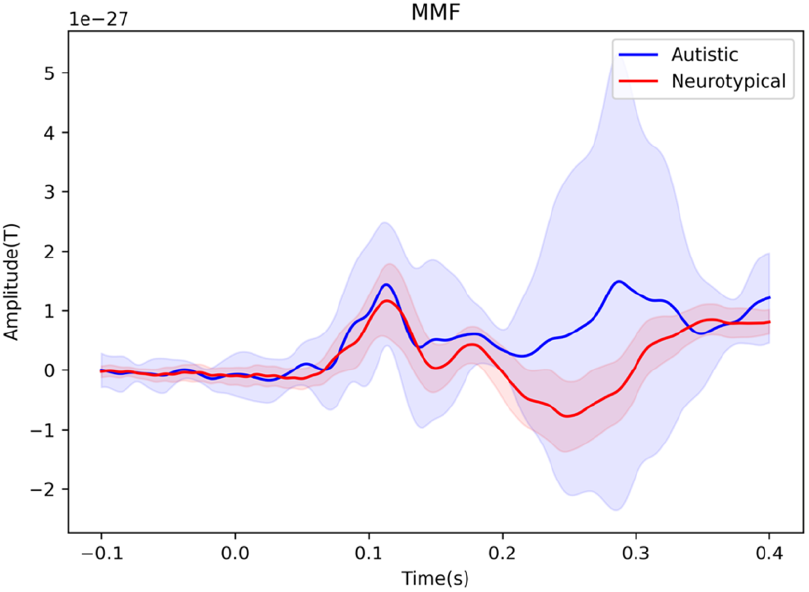
Group-level global field power (GFP) MMF amplitudes (Tesla) by time (ms) for autistic children (blue waveform; *N* = 10) and neurotypical children (red waveform; *N* = 63).

### 3.1.2 Exploratory Case-Cohort Comparisons

Figure 4 shows the exploratory case-cohort analyses, comparing each of the 10 autistic children to a group of average-age-matched neurotypical children. The leftmost panels in Figure 4 show each of the autistic children’s ‘MMF-GFP’ (red waveform) compared to that of their neurotypical counterparts’ (blue waveform). Consistent with the hypothesis derived from the HIPPEA predictive coding account of autism, six of the 10 autistic children (P2, P3, P5, P6, P8 and P9) showed significantly larger MMF amplitudes relative to their neurotypical counterparts, *p* < .05. The latencies of these significant clusters are summarised in Table 1 and shaded in red in Figure 4. Two of the autistic children,P1 and P4, likewise showed larger MMF amplitudes relative to neurotypical children, although these differences did not reach significance. Two other autistic children, P7 and P10, showed the opposite effect of larger standard-relative-to-deviant responses (i.e., ‘reverse MMF’ responses), *p* < .05.

**Figure 4.**
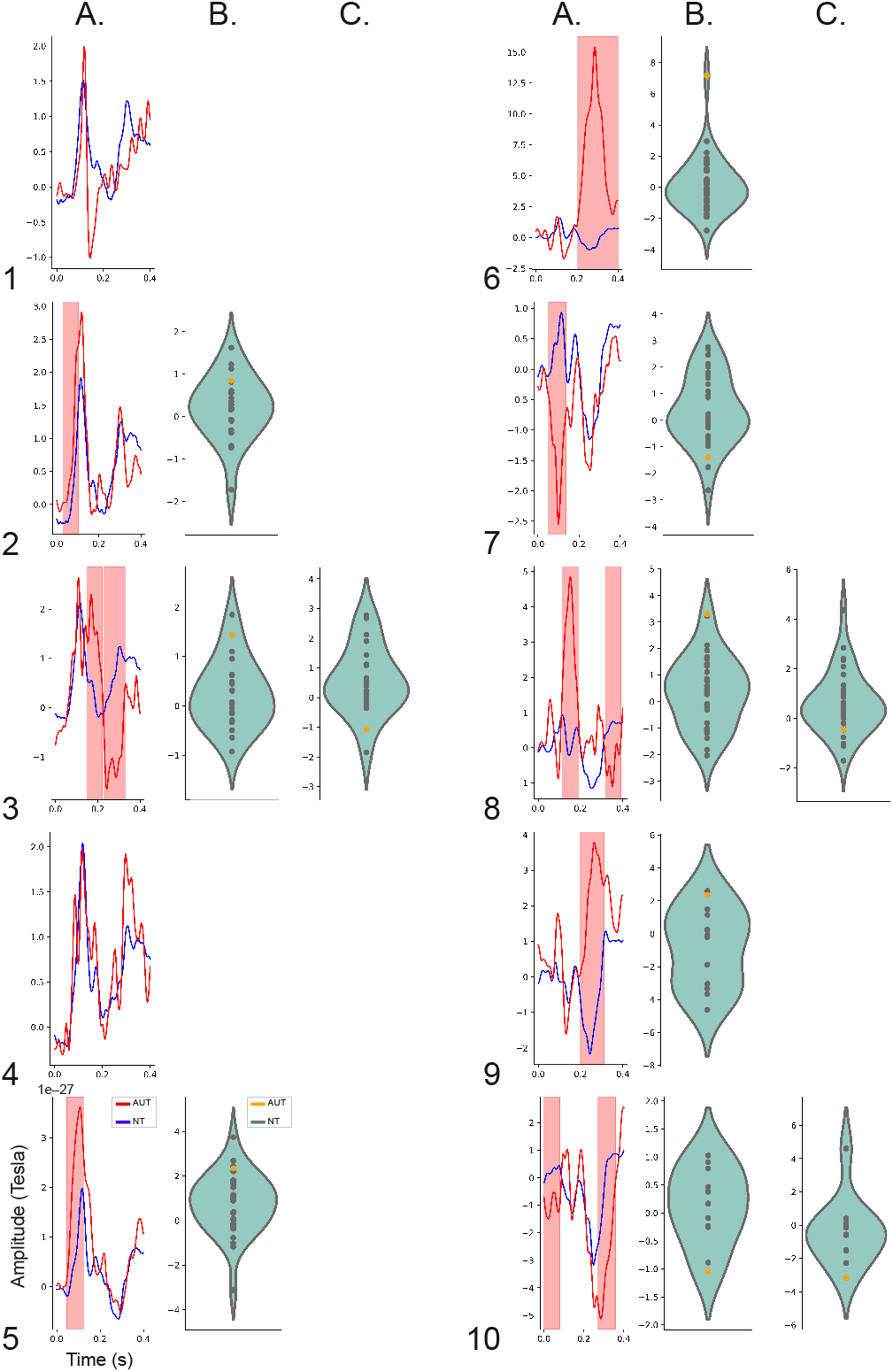
Results of the 10 case-cohort comparisons. Panel A: Global field power (GFP) MMF amplitudes (Tesla) by time (ms) for each individual autistic child (red waveforms), and their respective neurotypical counterparts (blue waveform). The shaded red areas indicate clusters of significant differences between the autistic child and the neurotypical group, *p* <.01. Panels B and C: For each significant cluster, we plotted the mean MMF amplitude (across the significant time-window) of the autistic child (yellow dot in the Violin plot) against each of the neurotypical average-age-matched children (green dots). Note the different y-axis scales.

## 4. Discussion

The aim of the current study was to evaluate several competing predictive coding accounts of autism. We hypothesised that attenuated MMF amplitudes in autistic children, compared to their neurotypical counterparts, would be consistent with the ‘inflexibly low prior-relative-to-sensory precision’ accounts (Brock, 2012; Pellicano, 2013; Pellicano & Burr, 2012). Furthermore, we hypothesised that relatively larger MMF amplitudes in autistic children would be consistent with the ‘HIPPEA’ account (Van de Cruys et al., 2013, 2014).

Unexpectedly, our group-level analysis yielded no significant difference in the magnitude of the MMF amplitude between the autistic and neurotypical children. We suspect that this null result may be partly due to an age-related confound, as both the autistic and the neurotypical groups were characterised by a wide range of ages (4.2–8.6 years and 3.0–9.8 years, respectively). Thus, any potential MMF amplitude differences between the autistic and neurotypical children may have been obscured by within-group maturational changes in the MMF waveform. This explanation may also account for the null results of several previous studies, which also compared MMN/F between groups of autistic and neurotypical children characterised by a wide range of ages (Gomot et al., 2002, 2011; Lepistö et al., 2009; Roberts et al., 2011; Weismuller et al., 2015).

To address this concern, we conducted a series of exploratory case-cohort analyses, comparing the MMF amplitudes of each of the autistic children to that of their average-age-matched neurotypical counterparts. Consistent with the hypothesis derived from the ‘HIPPEA’ account (Van de Cruys et al., 2013, 2014), six of the 10 autistic children showed significantly larger MMF amplitudes relative to their neurotypical counterparts (NB. an additional two autistic children, P1 and P4, likewise showed larger MMF amplitudes relative to neurotypical children, yet these differences did not reach significance). Under the HIPPEA account, the pseudo-random ‘deviants’ may have elicited overly precise PE signals, thereby motivating the brain to update its Bayesian model based on the unpredicted information. Thus, all subsequent repetitions of the deviant (which, through their repetition, became ‘standards’) may have been precisely predicted, thereby eliciting relatively minimal PE signals. Overall, this may have given rise to the relatively larger PE (i.e., evoked response) signals to the ‘deviants’ relative to the standards, resulting in relatively large MMF amplitudes in the autistic children.

Larger MMF amplitudes in autistic compared to neurotypical children is at odds with the results of a recent meta-analysis, which reported relatively attenuated MMN/F amplitudes in young autistic children (Schwartz et al., 2018). However, the children included in the meta-analysis were older (age range: 6 – 15 years) than those included in the current study (age range: 4.2–8.6 years). When we instead examined the findings from amongst the large number of individual studies, the current findings fell within the broad spectrum of these previous reports of autistic children showing MMN/F amplitudes that were smaller (Andersson et al., 2013; Dunn et al., 2008; Jansson-Verkasalo et al., 2005; Ludlow et al., 2014; Seri et al., 1999; Yoshimura et al., 2017), larger (Ferri et al., 2003), or not significantly different from that of their neurotypical counterparts (Gomot et al., 2002, 2011; Lepistö et al., 2009; Roberts et al., 2011; Weismuller et al., 2015).

The current findings were also at odds with the hypothesis derived from the account that autistic people have ‘inflexibly low prior-relative-to-sensory precision’ (Brock, 2012; Pellicano, 2013; Pellicano & Burr, 2012). However, the notion that autistic people have overly precise (Van de Cruys et al., 2013, 2014)—rather than overly weak priors—is arguably more consistent with autistic traits. For example, the high prevalence of savant skills among autistic people (Howlin et al., 2009) is more easily accounted for by a system that predicts the world with heightened precision, rather than a system that has imprecise expectations about the world (Van de Cruys et al., 2013). Furthermore, autistic preference for sameness (APA, 2013) may be better accounted for a system that maintains a high learning rate in otherwise volatile environments (an experience which may be cognitively taxing and overwhelming), rather than a system that maintains broad priors that account for environmental volatility.

This study has several limitations. The first of these relates to our small sample size. In general, it can be difficult to recruit autistic children for research participation due to challenges related to going to new places, working with new people, and breaking from familiar routines. Recruitment for the current study was made more challenging due to the 2020–21 COVID-19 lockdowns and restrictions. Furthermore, of the 21 autistic children who we were able to assess, 11 were excluded from the final analysis (see Supplementary Table S1), resulting in a total of 10 useable datasets (i.e., 48% of the original sample size). The most common reason for excluding participants was due to sensory aversions to the testing environment (especially having to wear the MEG cap, n = 7). Given our small sample size and the high percentage of participant exclusions (particularly of younger children with more severe language and sensory difficulties), we cannot be sure that the current findings are representative of the broader population of autistic children.

Second, although the current findings favour the HIPPEA account, it should be noted that not all the case-cohort comparisons were consistent with this account. Indeed, cases P7 and P10 showed the opposite effect of larger standard-relative-to-deviant responses (i.e., ‘reverse MMF’ responses). Although P7’s response fell within the neurotypical continuum (albeit at the extreme end of the spread), P10’s response was placed as the most extreme response for both significant time windows. Unfortunately, these divergent responses are difficult to explain as they do not fit neatly with our theory-driven hypotheses, nor with previous findings. Such divergence has not been observed in previous studies as past analyses have been constrained to group-level comparisons. Nevertheless, divergent findings were not unexpected, given the significant heterogeneity across the autism spectrum (Jeste & Geschwind, 2014). Investigating whether such divergent electrophysiological responses can be linked with autistic traits could be a fruitful topic for future research. Unfortunately, the sample size in the current study was too small to conduct such an investigation.

In conclusion, we found larger MMF amplitudes in most of the autistic children compared to their neurotypical average-age-matched counterparts. We interpret these findings as preliminary evidence in support of the ‘inflexibly high prior and sensory precision’ account, and against the ‘inflexibly low prior-relative-to-sensory precision’ accounts of autism. However, given the exploratory nature of the case-cohort analyses, as well as the small sample of autistic children on which it was based, the results need both to be replicated and modelled on a trial-by-trial basis in future work if they are to be used in strong support of the HIPPEA account.

## ACKNOWLEDGEMENTS

We are extremely grateful to all the children and families who so generously gave up their time to participate in this research.

## ETHICS STATEMENT

This research was conducted in accordance with the ethical guidelines of Macquarie

University. All protocols were approved by the Macquarie University Human Research

Ethics Committee (reference numbers: 5201300834 and 5201600188).

## CONSENT STATEMENT

Children and their parents provided verbal and written informed consent, respectively, before the experiment.

Hannah Rapaport^a,b^, Elizabeth Pellicano^b,c^, Robert A. Seymour^d^, Nick Benikos^a^, Wei He^a^, Yanan Sun^a^, Jon Brock^a^, Paul F. Sowman^a^

^a^School of Psychological Sciences, Macquarie University, Sydney, Australia

^b^Macquarie School of Education, Macquarie University, Sydney, Australia

^c^Department of Clinical, Educational and Health Psychology, University College London London, WC1E 6BT, United Kingdom

^d^Wellcome Centre for Human Neuroimaging, University College London, London, WC1N 3AR, United Kingdom

## CREDIT AUTHOR STATEMENT

**Hannah Rapaport**: Conceptualisation, Methodology, Software, Formal analysis, Investigation, Data Curation, Writing – Original Draft, Writing – Review & Editing, Visualization,**Elizabeth Pellicano**: Conceptualisation, Methodology, Writing – Review * Editing, Supervision, **Robert A. Seymour**: Methodology, Software, Writing – Review & Editing, Investigation, Visualization, **Nicholas Benikos:**Investigation, Resources, **Wei He**: Methodology, Investigation, Supervision, **Yanan Sun**: Software, **Jon Brock**: Conceptualisation, Writing – Review & Editing, Funding acquisition, **Paul F. Sowman**: Conceptualization, Methodology, Software, Formal analysis, Writing – Review & Editing, Visualization, Supervision, Funding acquisition.

## Footnotes

1 Here the MMF was liberally defined as any significantly larger deviant-relative-to-standard waveform amplitude occurring within the 500 ms time window. This definition stands in contrast to the definition of the classic adult MMN/F, which emerges soon after the second ‘N1 or ‘M2’ component (Hari & Puce, 2017, p. 265; Näätänen et al., 2019a, p. 53) and is visible between 100 to 250 ms following stimulus onset (Näätänen et al., 2019b). Our reason for relying on this more liberal definition is that the latency of the peak mismatch effect changes across childhood (Näätänen et al., 2019a). As such, it would have been inappropriate to constrain our analysis to the classic adult MMN time window. It should be noted that earlier paediatric auditory oddball studies have likewise used liberally-defined time windows to extract MMN effects (e.g., 200-330 ms in 6-to 7-year-olds, Lovio et al., 2009; 300-550 ms in 4-to 12-year-olds, Partanen et al., 2013; 150-400 ms in 5-to 7-year-olds, Petermann et al., 2009; 100-300 ms in 9-to 13-year-olds, Putkinen et al., 2014; 100-320 ms in 4-to 10-year-olds, Shafer et al., 2000).

## References

American Psychiatric Association. (2013). Diagnostic and statistical manual of mental disorders (5th ed.). Washington, DC. https://doi.org/https://doi.org/10.1176/appi.books.9780890425596

Andersson, S., Posserud, M.-B. B., & Lundervold, A. J. (2013). Early and late auditory event-related potentials in cognitively high functioning male adolescents with autism spectrum disorder. Research in Autism Spectrum Disorders, 7(7), 815–823. https://doi.org/10.1016/j.rasd.2013.03.007

Brock, J. (2012). Alternative Bayesian accounts of autistic perception: Comment on Pellicano and Burr. Trends in Cognitive Sciences, 16(12), 573–574. https://doi.org/10.1016/j.tics.2012.10.005

Centers for Disease Control and Prevention. (2021). Preschooler (3-5 years old). U.S. Department of Health & Human Services. https://www.cdc.gov/ncbddd/childdevelopment/positiveparenting/preschoolers.html

Clark, A. (2013). Whatever next? Predictive brains, situated agents, and the future of cognitive science. Behavioral and Brain Sciences, 36(3), 181–204. https://doi.org/10.1017/S0140525X12000477

Clark, A. (2015). Surfing uncertainty: Prediction, action, and the embodied mind. Oxford University Press. https://doi.org/10.1093/acprof:oso/9780190217013.001.0001

Corsello, C., Hus, V., Pickles, A., Risi, S., Cook, E. H., Leventhal, B. L., & Lord, C. (2007). Between a ROC and a hard place: Decision making and making decisions about using the SCQ. Journal of Child Psychology and Psychiatry, 48(9), 932–940. https://doi.org/10.1111/J.1469-7610.2007.01762.X

de Cheveigné, A., & Simon, J. Z. (2007). Denoising based on time-shift PCA. Journal of Neuroscience Methods, 165(2), 297–305. https://doi.org/10.1016/J.JNEUMETH.2007.06.003

Denham, S. L., & Winkler, I. (2020). Predictive coding in auditory perception: Challenges and unresolved questions. The European Journal of Neuroscience, 51(5), 1151–1160. https://doi.org/10.1111/ejn.13802

Dunn, M. A., Gomes, H., & Gravel, J. (2008). Mismatch negativity in children with autism and typical development. Journal of Autism and Developmental Disorders, 38(1), 52–71. https://doi.org/http://dx.doi.org/10.1007/s10803-007-0359-3

Ferri, R., Elia, M., Agarwal, N., Lanuzza, B., Musumeci, S. A., & Pennisi, G. (2003). The mismatch negativity and the P3a components of the auditory event-related potentials in autistic low-functioning subjects. Clinical Neurophysiology, 114(9), 1671–1680. https://doi.org/10.1016/S1388-2457(03)00153-6

Friston, K. (2005). A theory of cortical responses. Philosophical Transactions of the Royal Society B: Biological Sciences, 360(1456), 815–836. https://doi.org/10.1098/rstb.2005.1622

Friston, K., Lawson, R., & Frith, C. D. (2013). On hyperpriors and hypopriors: Comment on Pellicano and Burr. Trends in Cognitive Sciences, 17(1), 1. https://doi.org/10.1016/j.tics.2012.11.003

Frith, U. (1989). Au/ism: Explaining the enigma (Vol. 3). Oxford: Blackwell Publishing.

Frith, U., & Happé, F. (1994). Autism: Beyond “theory of mind.” Cogni/ion, 50(1-3), 115–132. https://doi.org/10.1016/0010-0277(94)90024-8

Garrido, M. I., Friston, K. J., Kiebel, S. J., Stephan, K. E., Baldeweg, T., & Kilner, J. M. (2008). The functional anatomy of the MMN: A DCM study of the roving paradigm. NeuroImage, 42(2), 936–944. https://doi.org/10.1016/J.NEUROIMAGE.2008.05.018

Gibbs, V., Aldridge, F., Sburlati, E., Chandler, F., Smith, K., & Cheng, L. (2019). Missed opportunities: An investigation of pathways to autism diagnosis in Australia. Research in Autism Spectrum Disorders, 57, 55–62. https://doi.org/10.1016/j.rasd.2018.10.007

Gomot, M., Blanc, R., Clery, H., Roux, S., Barthelemy, C., & Bruneau, M. (2011). Candidate electrophysiological endophenotypes of hyper-reactivity to change in autism. Journal of Autism and Developmental Disorders, 41(6), 705–714. https://doi.org/http://dx.doi.org/10.1007/s10803-010-1091-y

Gomot, M., Giard, M.-H., Adrien, J.-L., Barthelemy, C., & Bruneau, N. (2002). Hypersensitivity to acoustic change in children with autism: Electrophysiological evidence of left frontal cortex dysfunctioning. Psychophysiology, 39(5), 577–584. https://doi.org/10.1111/1469-8986.3950577

Gotham, K., Pickles, A., & Lord, C. (2009). Standardising ADOS scores for a measure of severity in autism spectrum disorders. Journal of Autism and Developmental Disorders, 39(5), 693–705. https://doi.org/10.1007/s10803-008-0674-3

Gramfort, A., Luessi, M., Larson, E., Engemann, D. A., Strohmeier, D., Brodbeck, C., Goj, R., Jas, M., Brooks, T., Parkkonen, L., & Hämäläinen, M. (2013). MEG and EEG data analysis with MNE-Python. Frontiers in Neuroscience, 7, 267. https://doi.org/10.3389/fnins.2013.00267

Happé, F., & Booth, R. D. L. (2008). The power of the positive: Revisiting weak coherence in autism spectrum disorders. Quarterly Journal of Experimental Psychology, 61(1), 50–63. https://doi.org/10.1080/17470210701508731

Happé, F., & Frith, U. (2006). The weak coherence account: Detail-focused cognitive style in autism spectrum disorders. Journal of Autism and Developmental Disorders, 36(1), 5–25. https://doi.org/10.1007/s10803-005-0039-0

He, W., Donoghue, T., Sowman, P. F., Seymour, R. A., Brock, J., Crain, S., Voytek, B., & Hillebrand, A. (2019). Co-increasing neuronal noise and beta power in the developing brain. BioRxiv, 839258. https://doi.org/10.1101/839258

Heilbron, M., & Chait, M. (2017). Great expectations: Is there evidence for predictive coding in auditory cortex? Neuroscience, 389, 54–73. https://doi.org/10.1016/J.NEUROSCIENCE.2017.07.061

Hohwy, J. (2013). The predictive mind. Oxford University Press. https://doi.org/10.1093/acprof:oso/9780199682737.001.0001

Howlin, P., Goode, S., Hutton, J., & Rutter, M. (2009). Savant skills in autism: Psychometric approaches and parental reports. Philosophical Transactions of the Royal Society B: Biological Sciences, 364(1522). https://doi.org/10.1098/rstb.2008.0328

Jansson-Verkasalo, E., Ceponiene, R., Kielinen, M., Suominen, K., Jäntti, V., Linna, S. L., Moilanen, I., & Näätänen, R. (2003). Deficient auditory processing in children with Asperger Syndrome, as indexed by event-related potentials. Neuroscience Letters, 338(3), 197–200. https://doi.org/10.1016/S0304-3940(02)01405-2

Jansson-Verkasalo, E., Kujala, T., Jussila, K., Mattila, M. L., Moilanen, I., Näätänen, R., Suominen, K., & Korpilahti, P. (2005). Similarities in the phenotype of the auditory neural substrate in children with Asperger syndrome and their parents. European Journal of Neuroscience, 22(4), 986–990. https://doi.org/10.1111/j.1460-9568.2005.04216.x

Jeste, S. S., & Geschwind, D. H. (2014). Disentangling the heterogeneity of autism spectrum disorder through genetic findings. Nature Reviews Neurology, 10(2), 74–81. https://doi.org/10.1038/nrneurol.2013.278

Johnson, B. W., Crain, S., Thornton, R., Tesan, G., & Reid, M. (2010). Measurement of brain function in pre-school children using a custom sized whole-head MEG sensor array. Clinical Neurophysiology, 121(3), 340–349. https://doi.org/10.1016/j.clinph.2009.10.017

Knösche, T. R. (2002). Transformation of whole-head MEG recordings between different sensor positions/Transformation von Ganzkopf-MEG-Messungen zwischen verschiedenen Sensorpositionen. Biomedizinische Technik, 47(3), 59–62. https://doi.org/10.1515/bmte.2002.47.3.59

Korpilahti, P., Jansson-Verkasalo, E., Mattila, M. L., Kuusikko, S., Suominen, K., Rytky, S., Pauls, D. L., & Moilanen, I. (2007). Processing of affective speech prosody is impaired in Asperger syndrome. Journal of Autism and Developmental Disorders, 37(8), 1539–1549. https://doi.org/10.1007/s10803-006-0271-2

Kujala, T., Kuuluvainen, S., Saalasti, S., Jansson-Verkasalo, E., von Wendt, L., Lepisto, T., Lepistö, T., Wendt, L. von, & Lepistö, T. (2010). Speech-feature discrimination in children with Asperger syndrome as determined with the multi-feature mismatch negativity paradigm. Clinical Neurophysiology, 121(9), 1410–1419. https://doi.org/http://dx.doi.org/10.1016/j.clinph.2010.03.017

Lawson, R. P., Mathys, C., & Rees, G. (2017). Adults with autism overestimate the volatility of the sensory environment. Nature Neuroscience, 20(9), 1293–1299. https://doi.org/10.1038/nn.4615

Lawson, R. P., Rees, G., & Friston, K. J. (2014). An aberrant precision account of autism. Frontiers in Human Neuroscience, 8, 302. https://doi.org/10.3389/fnhum.2014.00302

Le Couteur, A., Rutter, M., Lord, C., Rios, P., Robertson, S., Holdgrafer, M., & McLennan, J. (1989). Autism diagnostic interview: A standardised investigator-based instrument. Journal of Autism and Developmental Disorders 1989 19:3, 19(3), 363–387. https://doi.org/10.1007/BF02212936

Lepistö, T., Kajander, M., Vanhala, R., Alku, P., Huotilainen, M., Näätänen, R., & Kujala, T. (2008). The perception of invariant speech features in children with autism. Biological Psychology, 77(1), 25–31. https://doi.org/http://dx.doi.org/10.1016/j.biopsycho.2007.08.010

Lepistö, T., Kuitunen, A., Sussman, E., Saalasti, S., Jansson-Verkasalo, E., Nieminen-von Wendt, T., & Kujala, T. (2009). Auditory stream segregation in children with Asperger syndrome. Biological Psychology, 82(3), 301–307. https://doi.org/10.1016/j.biopsycho.2009.09.004

Lepistö, T., Kujala, T., Vanhala, R., Alku, P., Huotilainen, M., & Näätänen, R. (2005). The discrimination of and orienting to speech and non-speech sounds in children with autism. Brain Research, 1066(1–2), 147–157. https://doi.org/http://dx.doi.org/10.1016/j.brainres.2005.10.052

Lepistö, T., Silokallio, S., Nieminen-von Wendt, T., Alku, P., Näätänen, R., Kujala, T., Naatanen, R., & Kujala, T. (2006). Auditory perception and attention as reflected by the brain event-related potentials in children with Asperger syndrome. Clinical Neurophysiology, 117(10), 2161–2171. https://doi.org/http://dx.doi.org/10.1016/j.clinph.2006.06.709

Lord, C., Brugha, T. S., Charman, T., Cusack, J., Dumas, G., Frazier, T., Jones, E. J. H., Jones, R. M., Pickles, A., State, M. W., Taylor, J. L., & Veenstra-VanderWeele, J. (2020). Autism spectrum disorder. Nature Reviews Disease Primers, 6(1), 1–23. https://doi.org/10.1038/s41572-019-0138-4

Lord, C., Rutter, M., DiLavore, P. C., Risi, S., Gotham, K., & Bishop, S. L. (2012). Autism diagnostic observation schedule (2nd ed.). Western Psychological Services.

Ludlow, A., Mohr, B., Whitmore, A., Garagnani, M., Pulvermüller, F., & Gutierrez, R. (2014). Auditory processing and sensory behaviours in children with autism spectrum disorders as revealed by mismatch negativity. Brain and Cognition, 86(1), 55–63. https://doi.org/10.1016/j.bandc.2014.01.016

Maris, E., & Oostenveld, R. (2007). Nonparametric statistical testing of EEG-and MEG-data. Journal of Neuroscience Methods, 164(1), 177–190. https://doi.org/10.1016/J.JNEUMETH.2007.03.024

Medvedovsky, M., Taulu, S., Bikmullina, R., & Paetau, R. (2007). Artifact issue during head position correction in MEG. Epilepsia, 48. https://oce.ovid.com/article/00003606-200710001-00567

Mottron, L., & Burrack, J. A. (2001). Enhanced perceptual functioning in the development of autism. In The development of autism: Perspectives from theory and research (pp. 131–148). Lawrence Erlbaum Associates Publishers. https://psycnet.apa.org/record/2001-01233-007

Mottron, L., Dawson, M., Soulières, I., Hubert, B., & Burack, J. (2006). Enhanced perceptual functioning in autism: An update, and eight principles of autistic perception. Journal of Autism and Developmental Disorders, 36(1), 27–43. https://doi.org/10.1007/s10803-005-0040-7

Näätänen, R., Kujala, T., & Light, G. (2019). The mismatch negativity (MMN): An introduction. In Mismatch negativity: A window to the brain (pp. 1–40). Oxford University Press. https://doi.org/10.1093/oso/9780198705079.003.0001

Oostenveld, R., Fries, P., Maris, E., & Schoffelen, J. M. (2011). FieldTrip: Open source software for advanced analysis of MEG, EEG, and invasive electrophysiological data. Computational intelligence and Neuroscience, 2011, 1–9. https://doi.org/10.1155/2011/156869

Oyama, D., Adachi, Y., Higuchi, M., Kawai, J., Kobayashi, K., & Uehara, G. (2012). Real-time coil position monitoring system for biomagnetic measurements. Physics Procedia, 36, 280–285. https://doi.org/10.1016/J.PHPRO.2012.06.160

Pellicano, E. (2013). Sensory symptoms in autism: A blooming, buzzing confusion? Child Development Perspectives, 7(3), 143–148. https://doi.org/10.1111/cdep.12031

Pellicano, E., & Burr, D. (2012). When the world becomes “too real”: A Bayesian explanation of autistic perception. Trends in Cognitive Sciences, 16(10), 504–510. https://doi.org/10.1016/j.tics.2012.08.009

Rao, R. P. N., & Ballard, D. H. (1999). Predictive coding in the visual cortex: A functional interpretation of some extra-classical receptive-field effects. Nature Neuroscience, 2(1), 79–87. https://doi.org/10.1038/4580

Rapaport, H., Seymour, R. A., Sowman, P. F., Benikos, N., Stylianou, E., Johnson, B. W., Crain, S., & He, W. (2019). Studying brain function in children using magnetoencephalography. Journal of Visualized Experiments, 146, e58909. https://doi.org/10.3791/58909

Roberts, T. P. L., Cannon, K. M., Tavabi, K., Blaskey, L., Khan, S. Y., Monroe, J. F., Qasmieh, S., Levy, S. E., & Edgar, J. C. (2011). Auditory magnetic mismatch field latency: A biomarker for language impairment in autism. Biological Psychiatry, 70(3), 263–269. https://doi.org/10.1016/j.biopsych.2011.01.015

Rutter, M., Bailey, A., & Lord, C. (2003). The social communication questionnaire. Western Psychological Services.

Schwartz, S., Shinn-Cunningham, B., & Tager-Flusberg, H. (2018). Meta-analysis and systematic review of the literature characterising auditory mismatch negativity in individuals with autism. Neuroscience & Biobehavioral Reviews, 87, 106–117. https://doi.org/10.1016/J.NEUBIOREV.2018.01.008

Seri, S., Cerquiglini, A., Pisani, F., & Curatolo, P. (1999). Autism in tuberous sclerosis: Evoked potential evidence for a deficit in auditory sensory processing. Clinical Neurophysiology, 110(10), 1825–1830. https://doi.org/http://dx.doi.org/10.1016/S1388-2457%2899%2900137-6

Seymour, R. A. (2019). Macquarie-MEG-Research/MQ_MEG Scripts: v0.1 for Zenodo (Version v0.1zenodo). Zenodo. https://doi.org/10.5281/zenodo.3406897

Simmons, D. R., Robertson, A. E., McKay, L. S., Toal, E., McAleer, P., & Pollick, F. E. (2009). Vision in autism spectrum disorders. Vision Research, 49(22), 2705–2739. https://doi.org/10.1016/j.visres.2009.08.005

van Boxtel, J. J. A., & Lu, H. (2013). A predictive coding perspective on autism spectrum disorders. Frontiers in Psychology, 4, 19. https://doi.org/10.3389/fpsyg.2013.00019

Van de Cruys, S., Evers, K., Boets, B., Wagemans, J., De-Wit, L., Evers, K., Boets, B., & Wagemans, J. (2013). Weak priors versus overfitting of predictions in autism: Reply to Pellicano and Burr. I-Perception, 4(2), 95–97. https://doi.org/10.1068/i0580ic

Van de Cruys, S., Evers, K., Van der Hallen, R., van Eylen, L., Boets, B., De-Wit, L., & Wagemans, J. (2014). Precise minds in uncertain worlds: Predictive coding in autism. Psychological Review, 121(4), 649–675. https://doi.org/10.1037/a0037665

Weismuller, B., Thienel, R., Youlden, A.-M. M., Fulham, R., Koch, M., & Schall, U. (2015). Psychophysiological correlates of developmental changes in healthy and autistic boys. Journal of Autism and Developmental Disorders, 45(7), 2168–2175. https://doi.org/10.1007/s10803-015-2385-x

Winkler, I., Debener, S., Muller, K.-R., & Tangermann, M. (2015). On the influence of high-pass filtering on ICA-based artifact reduction in EEG-ERP. 2015 37th Annual International Conference of the IEEE Engineering in Medicine and Biology Society (EMBC), 4101–4105. https://doi.org/10.1109/EMBC.2015.7319296

Yoshimura, Y., Kikuchi, M., Hayashi, N., Hiraishi, H., Hasegawa, C., Takahashi, T., Oi, M., Remijn, G. B., Ikeda, T., Saito, D. N., Kumazaki, H., & Minabe, Y. (2017). Altered human voice processing in the frontal cortex and a developmental language delay in 3-to 5-year-old children with autism spectrum disorder. Scientific Reports, 7(1), 17116. https://doi.org/10.1038/s41598-017-17058-x

Yu, L., Fan, Y., Deng, Z., Huang, D., Wang, S., & Zhang, Y. (2015). Pitch processing in tonal-language-speaking children with autism: An event-related potential study. Journal of Autism and Developmental Disorders, 45(11), 3656–3667. https://doi.org/http://dx.doi.org/10.1007/s10803-015-2510-x

