## Supplemental Table S1 for "Hearing the World Differently: Examining Predictive Coding Accounts of Autism Using MEG"

**Supplementary Materials**

**Table S1**

Demographic Details of the Autistic Participants Excluded from the Analysis

| ID | Age (years) | Sex | SCQ | ADOS-2 | | | Reason for exclusion |
|---|---|---|---|---|---|---|---|
| | | | | CSS | Module | Item A1 | |
| 1 | 3.0 | M | 25 | 7 | 1 | Occasional phrases only; mostly single words | Sensory aversion to the MEG cap |
| 2 | 3.1 | M | 16 | 6 | 1 | Occasional phrases only; mostly single words | Sensory aversion to the MEG cap |
| 3 | 3.4 | M | 4 | 4 | 2 | Complex (i.e., Highest rating) | Sensory aversion to the auditory oddball stimuli |
| 4 | 3.6 | F | - | 9 | 1 | Fewer than five words used during the ADOS-2 evaluation | Sensory aversion to the MEG cap |
| 5 | 3.8 | M | 16 | 3 | 1 | Complex | Sensory aversion to the MEG cap |
| 6 | 3.9 | F | 15 | 9 | 1 | Fewer than five words used during the ADOS-2 evaluation | Sensory aversion to the MEG cap |
| 7 | 5.0 | M | 20 | 9 | 1 | Occasional phrases only; mostly single words | Sensory aversion to the digitiser pen. Would also not stay still long enough to perform the iPad digitisation. |
| 8 | 5.7 | M | 16 | 6 | 3 | Complex | Anxiety related to putting their head inside the MEG dewar |
| 9 | 5.9 | F | - | 5 | 1 | Complex | Anxiety related to putting their head inside the MEG dewar |
| 10 | 7.8 | M | 27 | 9 | 1 | No words or word approximations used meaningfully | Sensory aversion to the MEG cap |
| 11 | 8.3 | M | 23 | 8 | 3 | Complex | Head was too large for the child MEG dewar |

*Note.* SCQ = Social Communication Questionnaire (SCQ; Rutter et al., 2003) scores. ADOS-2 = Autism Diagnostic Observation Schedule 2 (ADOS-2; Lord et al., 2012). CSS = Calibrated Severity Score (Gotham et al., 2009). Item A1 = Overall Level of Non-Echoed Spoken Language.
